# Red muscle activity in bluegill sunfish *Lepomis macrochirus* during forward accelerations

**DOI:** 10.1101/459990

**Authors:** Margot A. B. Schwalbe, Alexandra L. Boden, Tyler N. Wise, Eric D. Tytell

**Affiliations:** Department of Biology, Tufts University, 200 Boston Ave, Ste 4700, Medford, MA 02155

**Keywords:** Locomotion, Swimming, EMG, co-activation, duty cycle

## Abstract

Fishes generate force to swim by activating muscles on either side of their flexible bodies. To accelerate, they must produce higher muscle forces, which leads to higher reaction forces back on their bodies from the environment. If their bodies are too flexible, the forces during acceleration cannot be transmitted effectively to the environment. Here, we investigate whether fish can use their red muscle to stiffen their bodies during acceleration. We used high-speed video, electromyographic recordings, and a new digital inertial measurement unit to quantify body kinematics, red muscle activity, and 3D orientation and centre of mass acceleration during forward accelerations and steady swimming over several speeds. During acceleration, fish co-activated anterior muscle on the left and right side, and activated all muscle sooner and kept it active for a larger fraction of the tail beat cycle. These activity patterns are consistent with our hypothesis that fish use their red muscle to stiffen their bodies during acceleration. We suggest that during impulsive movements, flexible organisms like fishes can use their muscles not only to generate propulsive power but to tune the effective mechanical properties of their bodies, increasing performance during rapid movements and maintaining flexibility for slow, steady movements.

## Introduction

Fishes swim by bending their bodies from side to side, but this undulatory motion is energetically expensive^1–3^. If a fish’s body is stiffer, it takes more energy to bend it. Fish with more flexible bodies should therefore use less energy to swim steadily, if all other factors are the same^4,5^. Yet, even fishes specialized for steady, long distance travel must also rapidly accelerate at times to escape predators or to capture prey. The body should be stiffer for rapid, impulsive movements so that the animal can effectively transmit large forces to the environment^4,6,7^. Many fishes must switch between steady swimming, where energy conservation would tend to favour more flexible bodies, and rapid, unsteady movements, where high performance would favour stiffer bodies. How are they able to do so? Here, we test the hypothesis, first posed by Blight^8^, that fishes may use their muscles to alter the effective stiffness of their bodies to manage these conflicting demands.

There is some evidence that fish can change their effective stiffness. In other animals, co-activation of groups of antagonist muscles increases the stiffness of a joint ^7,10,11^, and activation of individual muscles during lengthening, termed eccentric activation, also increases joint stiffness^12^. Very flexible fishes, such as lamprey and eels, may be able to increase the effective stiffness of their bodies by shifting the timing and duration of muscle activity ^6,7^. Less flexible fishes, such as trout, shift the timing of muscle activity during rapid, unsteady swimming ^9^, which may also increase relative body stiffness. Even moderately stiff fish like sunfishes and very stiff fish like tunas face the same trade-off between steady swimming performance and rapid acceleration.

Muscles, of course, also power movement. To swim, fish activate the axial muscles on either side of their bodies ^13^. These muscles generate forces that create a propulsive wave that travels the length of the fish’s body from head to tail, which in turn generates power that is converted to forward thrust by the tail ^14–16^. Most fishes have axial muscle with two distinct muscle fibre types: red, slow oxidative muscle that is found in a narrow wedge running longitudinally along the body just beneath the skin; and white, fast glycolytic muscle that is arranged in serial, nested cones, which form the majority of the musculature of the fish. Red muscle is active during steady, slow- to medium-speed swimming, and both red and white muscle are active during vigorous, high-speed unsteady swimming^17,18^.

As swimming speed increases, fish primarily increase their tail beat frequency, while tail amplitude stays relatively constant ^19^. To achieve the higher frequency movements, fish activate their muscles earlier and decrease how long their muscles are active in a tail beat ^13,18^. The duration of the muscle activity relative to the tail beat cycle generally decreases along the body, but depends on swimming speed, muscle type, and locomotion mode ^18,20^.

These patterns have been observed during steady swimming; however, much less is known about the neuromuscular control of unsteady swimming. What we do know primarily concerns C-start escape responses, which are rapid turning accelerations^21^. Routine linear accelerations, which are very common in nature^22^, have been studied much less and we know very little about the muscle activity that drives these behaviours. Unlike burst and glide swimming, they are characterized by relatively symmetrical tail beats at higher frequency and greater amplitude than steady swimming. Tytell^23^ found that eels increase both tailbeat frequency and amplitude during acceleration. Akanyeti et al.^24^ surveyed a large number of fish species during accelerations and found that this increase in both frequency and amplitude is very consistent across species with different body shapes and swimming modes. Ellerby and Altringham^9^ studied muscle function in sprinting rainbow trout and observed that white muscle turned on earlier at higher speeds, with more eccentric activity along the length of the body, potentially stiffening the body ^25^.

To our knowledge, no one has extensively studied the kinematics and motor patterns of red muscle during forward accelerations. Red muscle is active during rapid behaviours, including escape responses, even though it may not contract fast enough to contribute much power for the behaviour ^9,17^. Instead, we hypothesize that, along with generating power, it may function to tune the effective mechanical properties of the body. Even if it contracts too slowly to contribute much power for rapid acceleration, co-activation or eccentric activation of red muscle may help to stiffen the body and increase acceleration performance.

Studying how fish perform impulsive movements will advance the understanding of the locomotor movement itself, neuromuscular control, and the timing and distribution of forces during this behaviour. Therefore, to test the hypothesis that fish alter their effective body stiffness during forward accelerations, we examined kinematics and red muscle activity at different longitudinal positions in bluegill sunfish (*L. macrochirus*) during both steady swimming and forward accelerations over a range of swimming speeds. These measurements will not directly show changes in body stiffness, which is not possible to measure *in vivo*, but, together with what is known about how muscle stiffness varies depending on activation timing, they can demonstrate behaviour consistent with body stiffening. Increases in eccentric activation and co-activation as bluegill sunfish accelerate more rapidly would therefore indicate a mechanism for increasing effective body stiffness to improve force transmission for acceleration while maintaining performance during steady swimming. Such results will provide biologically inspired insights into the design and control of soft robots and actuators ^26^.

## Results

Bluegill sunfish altered their swimming kinematics and red muscle activity during accelerations compared to that during steady swimming over a range of speeds (1.5-2.5 body lengths per s, L s^-1^). Fig. 1 provides examples of raw data from a steady swimming trial (2.0 L s^-1^; Fig. 1A) and an acceleration trial (beginning at 1.5 L s^-1^; Fig. 1B). We recorded ventral video and used that to measure standard kinematic measurements including tail and head amplitude, tail beat frequency, body wavelength and wave speed. We used inertial measurement units to estimate the dynamic acceleration of the fish’s body (Figs. 1Aii, Bii), as well as its 3D orientation (Figs. 1Av, Bv). Finally, we recorded muscle activity at four locations in the axial red musculature, on both sides of the body (Figs. 1Aiv, Biv). During steady swimming, there is consistently a gap between muscle activity on opposite sides. During acceleration, EMG activity becomes more intense and the gap between activity on opposite sides becomes smaller, or activation will even overlap. Movies S1 and S2 show videos synchronized with the data shown in Fig. 1.

**Figure. 1.**
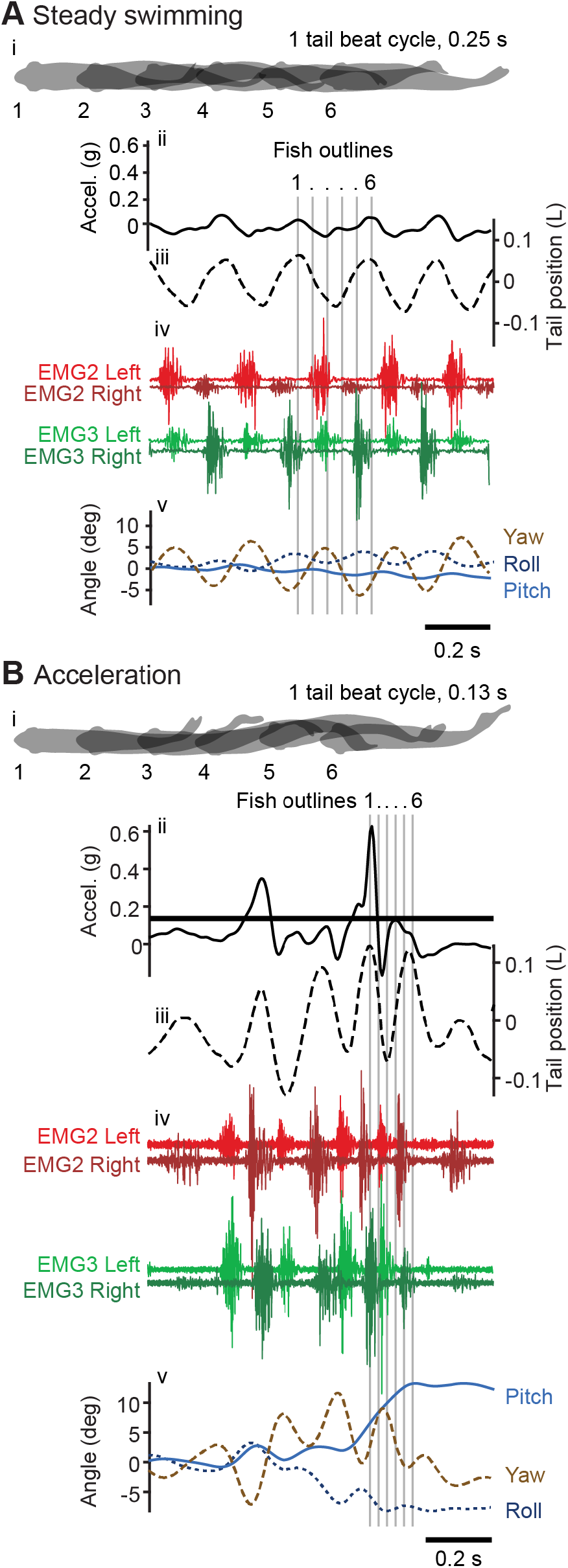
Examples of data from video, inertial measurement units, kinematic measurements, and electromyography. (A) Data collected from a steady swimming trial at 2.0 L s^-1^ and (B) an acceleration trial starting at 1.5 L s^-1^. Fish silhouettes at the top (i) are from one tail beat cycle from the representative trials (labelled 1-6) and are indicated by grey lines that extend through all traces. The panels below show (ii) forward acceleration (solid black line), (iii) the position of the tail (dash black line), (iv) EMG2 (red) and EMG 3 (green) from both left and right sides of the fish, and (v) roll (dark blue, dashed), pitch (blue), and yaw (brown, dashed). See Movies S1 and S2 for ventral video synchronized with the data shown in A and B, respectively.

Fig. 2 shows a summary of all of the kinematic data, including tail beat frequency, body wave speed, head and tail amplitude, body wavelength. We examined how all of these variables changed as a function of acceleration and swimming speed. We grouped swimming into steady and acceleration sequences and further characterized acceleration with three categories containing approximately the same number of tail beats: low (*a_dyn_* < 0.07*g*), medium (0.07 ≤ *a_dyn_* < 0.14 *g*), and high (*a_dyn_* ≥ 0.14*g*). The statistical results were robust to the number of categories. When bluegill sunfish accelerated, they significantly increased tail beat frequency, body wave speed, and head and tail amplitudes (Figs. 2A-D; p < 0.0001; Table S1). Body wavelength decreased significantly during acceleration (Fig. 2; p < 0.0001; Table S1). We found that tail beat frequency and tail amplitude increased significantly as acceleration increased from low to high (p < 0.0001, as indicated by the numbers above the acceleration categories in Fig. 2). Tail beat frequency and body wavelength increased significantly with swimming speed (p ≤ 0.0011) and body wave speed and head amplitude also tended to increase (p ≤ 0.0162), but tail amplitude did not change with swimming speed (p = 0.2165; Table S1).

**Figure 2.**
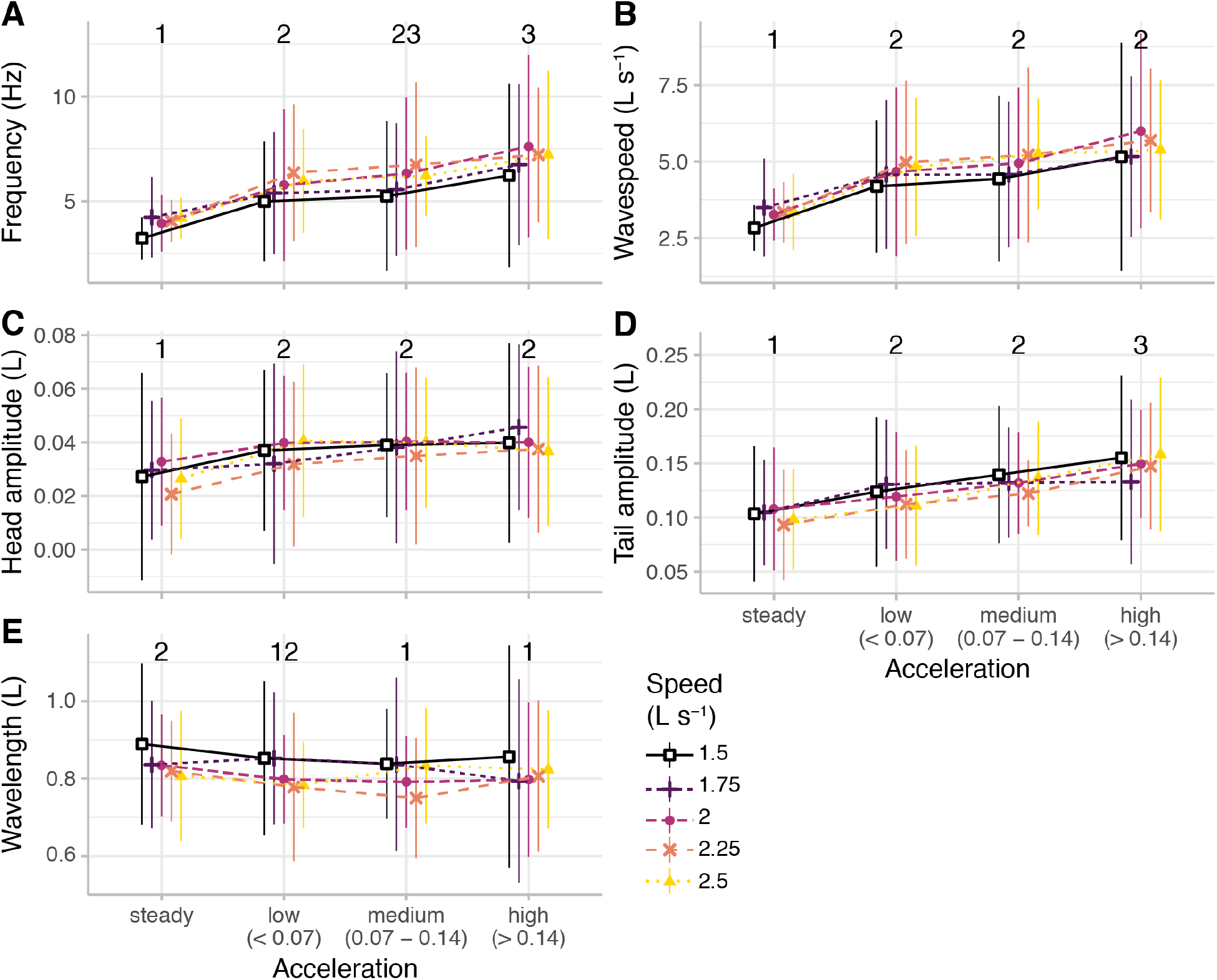
Bluegill sunfish (*N* = 5) altered swimming kinematics as acceleration increased. The plots show the relationship of acceleration and swimming speed with (A) tail beat frequency, (B) body wave speed, (C) head amplitude, (D) tail amplitude, (E) body wavelength, and (F) maximum muscle strain. Colour indicates the flow tank speed. Acceleration groups labelled with different numbers are significantly different from one another (p < 0.01). See Table S1 for statistical results.

To test the hypothesis that kinematics during accelerations are significantly different from those during steady swimming, we performed a principal components analysis (PCA) followed by a multivariate ANOVA (Fig. S1; Table S2). The PCA analysis included tail beat frequency, body wave speed, head and tail amplitudes, and body wavelength. The first two principal components explained 70.7% of the variation in the kinematic data. A Pillai trace indicated that kinematics were all significantly different across acceleration categories, swimming speeds, and their interaction (p < 0.0001 in all cases). The first principal component, which primarily included contributions from tail beat frequency, tail beat amplitude, and body wave speed, significantly differentiated acceleration categories (p < 0.0001) and speeds (p < 0.0001). The second component, which included contributions from head and body wavelength, also significantly differentiated swimming speeds (p < 0.0001), but not acceleration (p = 0.2766).

During accelerations, bluegill sunfish increase co-activation by activating red axial muscle longer relative to the tail beat cycle. Fig. 3 summarizes the muscle activity data. Burst duration decreased slightly during acceleration sequences (p < 0.0001; Table S3, Fig. 3A) and as swimming speed increased. However, the burst duration as a proportion of the tail beat cycle, called the duty cycle, did increase during accelerations (p < 0.0001; Table S2, Fig. 3B) because the tail beat frequency increased dramatically during acceleration (Fig. 2A), and did not change significantly with swimming speed (p = 0.3509). Both burst duration and duty cycle were significantly greater near the fish’s head (green colours in Fig. 3) and decreased along the body (p < 0.0001 in both cases; Figs. 3A,B). As duty cycle increases, it becomes more likely that muscle is active on both sides of the body simultaneously. We subsequently examined how long muscle activity overlapped at the same site on both sides of the body. A positive burst overlap indicates that left and right side muscles were active simultaneously, while a negative overlap indicates that there was a gap in time between opposite side activation. This burst overlap increased during accelerations (p *<* 0.0001; Table S3) but was lower in the posterior locations (p < 0.0001; Fig. 3C). Positive burst overlap, which indicates simultaneous activity across the left and right sides of the body, occurred more frequently during accelerations than during steady swimming most often at the anterior positions. At the highest accelerations (*a_dyn_* > 0.14*g*), 10.0% of bursts had positive burst overlaps, combining EMGs from all positions along the body.

**Figure 3.**
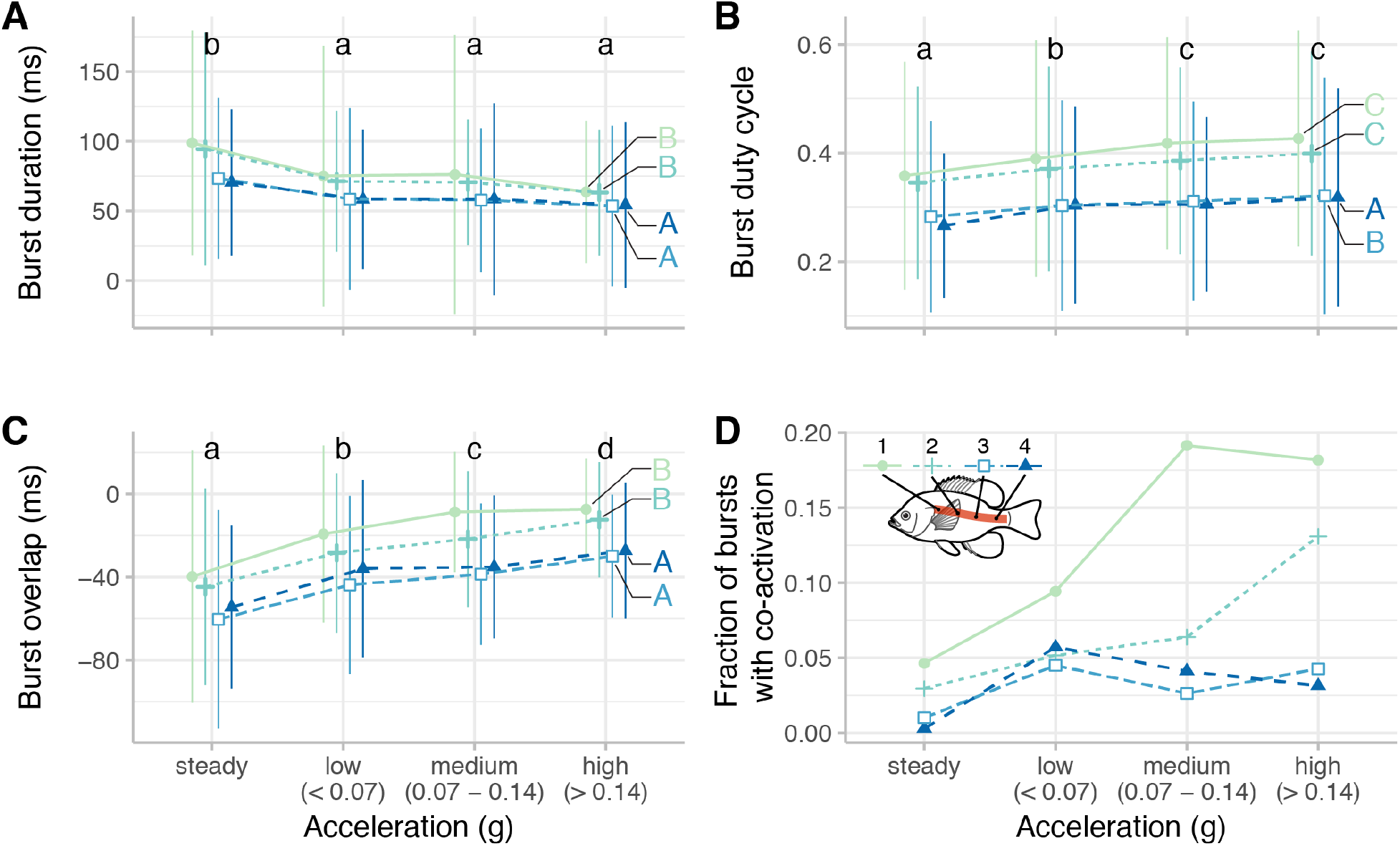
Bluegill sunfish (*N* = 5) alter their red axial muscle activity during forward accelerations. (A) Burst duration decreases during acceleration and along the length of the body. (B) Duty cycle increases with acceleration and is higher in the anterior portion of the body. (C) The duration of overlap between left and right side activity increases with accelerations and is higher in the anterior body. (D) The fraction of bursts with co-activation (overlap greater than zero) increases with acceleration, particularly for the most anterior location. Color indicates location along the body. Acceleration groups and EMG locations labeled with different lowercase or uppercase letters, respectively, are significantly different from one another (p < 0.01). See Table S2 for statistical results.

Bluegill sunfish also increase eccentric activation of red muscle, when the muscle is activated during lengthening. We compared the onset of muscle activity to body curvature at the same location as the EMG recording (Fig. 4A), computing a phase value that is zero at the beginning of muscle shortening. We performed a two-way test, equivalent to an ANOVA with circular data, called a Harrison-Kanji test^27^ to examine how burst onset and offset changed relative to acceleration and position along the body. During acceleration, burst onsets became significantly earlier (p < 0.0001; Table S4), but offsets did not change significantly (p = 0.0319). Bursts also started and ended earlier in more posterior locations (p < 0.0001 in both cases; Table S4). In absolute terms, during steady swimming, anterior muscle activated 5±24 ms before peak curvature, while posterior muscle activated 63±16 ms before peak curvature. At the highest acceleration, anterior muscle activated earlier (13±12 ms before peak curvature), while other muscle activated slightly later (25±10, 31±19, and 43±16 ms before peak curvature for EMGs 2, 3, and 4, respectively). At these high accelerations, posterior muscle is active later in absolute terms, but earlier in relative phase because the tail beat period is much shorter.

**Figure 4.**
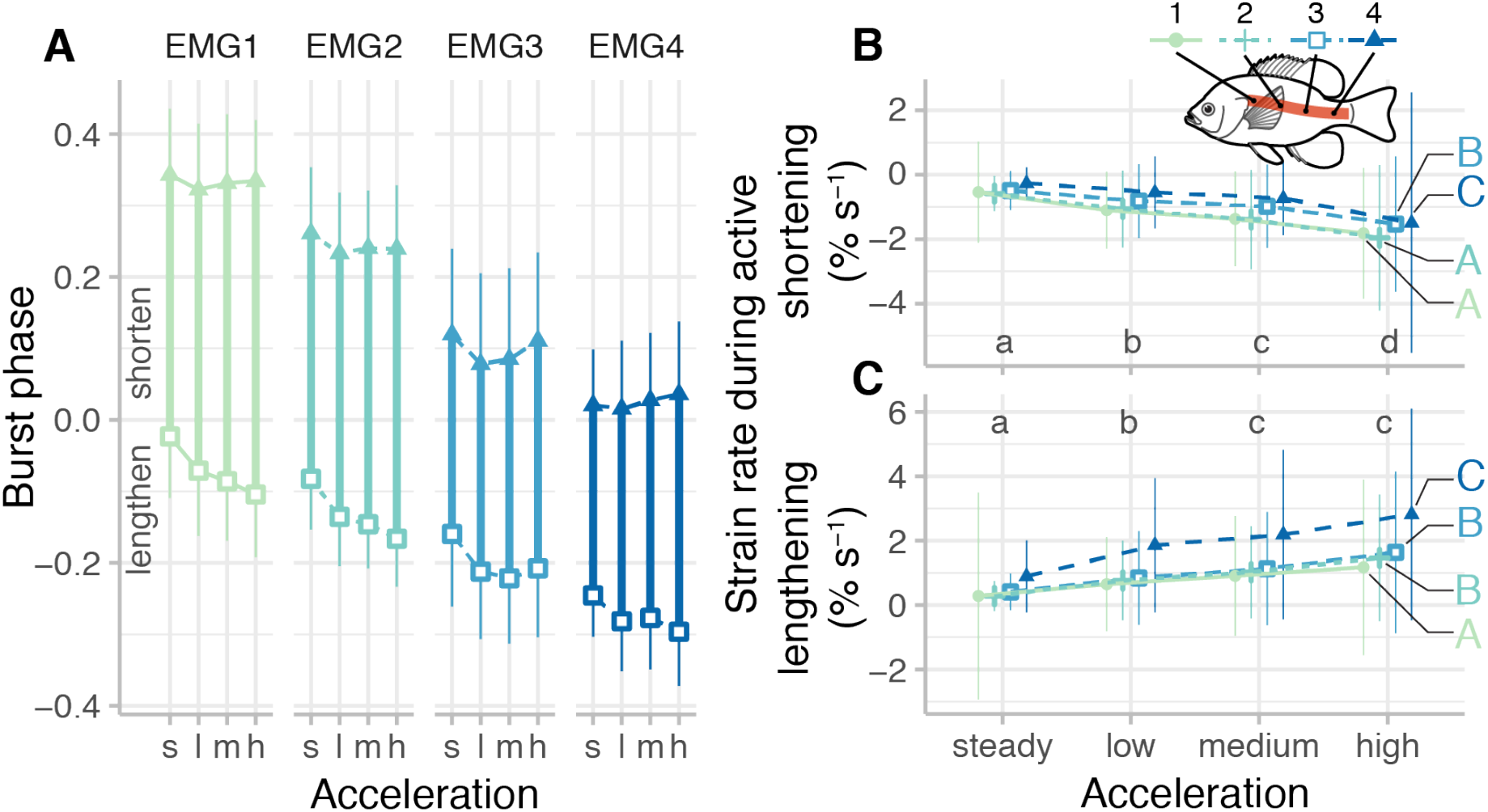
Bluegill sunfish (*N* = 5) increased eccentric contractions during forward accelerations versus steady swimming. (A) Onset and offset of muscle activation relative to body curvature, showing when the red muscle was active during lengthening (phase = -0.5-0) or shortening (phase = 0-0.5). See Table S3 for statistical results. (B) While the muscle was actively shortening, strain rate increased in magnitude (became more negative) with increasing acceleration but decreased along the body. (C) While the muscle was lengthening, strain rate increased with increasing acceleration and along the length of the body. Acceleration groups and EMG locations labeled with different lowercase or uppercase letters, respectively, are significantly different from one another (p < 0.01). See Table S2 for statistical results.

During accelerations, the strain rate during muscle activity, including both shortening and lengthening rates, increased in magnitude (Figs. 4B,C). Active muscle strain rates during acceleration became larger both during shortening and during lengthening (p < 0.0001; Table S3; Figs. 4B,C), reflecting the increase in overall strain and tail beat frequency during acceleration. In posterior muscle, the active lengthening rate became dramatically higher (blue colours in Fig. 4C).

## Discussion

The ability to manoeuvre effectively is essential for the survival of nearly all animals. Fish are capable of performing a variety of swimming manoeuvres with their flexible bodies, from slow, steady swimming to rapid escape responses. The kinematics and neuromuscular control of steady swimming and escape responses in fish are well-established ^21,28,29^, yet accelerations, an intermediate step between these swimming modes, have remained relatively unexplored (but see refs. ^23,24,30^). Previous computational work^4,5^ suggested that stiffer fish should accelerate more rapidly, if all other aspects of the body and swimming pattern are kept equal. We therefore hypothesized that fish use their slow-twitch red muscle to actively stiffen their bodies via bilateral co-activation and eccentric activation during accelerations. This would enable them to remain relatively flexible during steady swimming, which may reduce its energetic cost ^3–5^, but allow them to become effectively stiffer during high speed acceleration. Co-activation of antagonistic muscles and eccentric muscle activity are both known to increase effective stiffness across joints^31–33^, in isolated muscle preparations^34^, and in whole fishes^7,35^. We recorded the kinematics and red muscle activity patterns of bluegill sunfish, *Lepomis macrochirus*, during steady swimming over a range of speeds and during accelerations at different rates. Supporting our hypothesis, we found that the fish co-activated left and right side muscle (Figs. 3C,D) and activated them earlier in the cycle (Fig. 4A) as acceleration increased.

We found that tail beat frequency, body wave speed, and head and tail amplitude all increase significantly during acceleration compared to steady swimming (Fig. 2). In a recent study, Akanyeti et al.^24^ compared acceleration to steady swimming in a wide range of fish species. They found the same kinematic pattern that we observed, but they were not able to estimate the magnitude of the acceleration. Our study is the first to quantitatively compare how kinematics change relative to acceleration. In particular, we find that tail beat frequency and amplitude increase proportionally as bluegill sunfish accelerate faster. In experiments with flapping panels, the thrust output is proportional to the tail tip velocity, which is the product of frequency and amplitude ^36^. Thus, it seems clear that thrust is increasing as acceleration increases.

Overall, the kinematic pattern is affected both by swimming speed and by acceleration. During steady swimming, bluegill sunfish increased their tail beat frequency as the swimming speed increased (Fig. 2A) but did not change their tail amplitude (Fig. 2D), consistent with previous observations^16,19,29^. As acceleration increases, bluegill sunfish increase their tail beat frequency and body wave speed (Figs. 2A,B). In general, kinematics depend significantly on both swimming speed and acceleration. Kinematic variables for lower magnitude accelerations starting from higher speeds are different from those from a higher magnitude accelerations starting from a lower speed (Fig. S1; Table S2).

The muscle activity data are consistent with our hypothesis that fish actively stiffen their bodies during acceleration. During accelerations, bluegill sunfish activated their red muscle for a longer fraction of the tail beat cycle (Fig. 4B) and started the activity earlier during the lengthening phase compared to the activity during steady swimming (Fig. 4A). Previous studies have shown that duty cycle (the fraction of each tail beat that the muscle is active) decreases along the body during steady swimming and does not change substantially as speed increases ^14,15,37^, which was also the pattern observed here (Fig. 3B). During accelerations, duty cycle increased at each of the four EMG positions along the body. Higher duty cycle correlates with greater co-activation, as seen in the increase in positive burst overlap (Fig. 3D). During steady swimming only 2% of bursts had any positive overlap, but during all accelerations, 7.6% of the bursts had some amount of overlap. The increase in burst overlap was most pronounced at the most anterior EMG position, with 17% of muscle activity bursts at that location showing co-activation during medium and high accelerations.

Along with the increase in duty cycle, fish activated their red muscle earlier in the tail beat cycle during accelerations than during steady swimming. At anterior positions, muscle was active almost 8% of a tail beat cycle earlier during accelerations than during steady swimming (Fig. 4A). Ellerby and Altringham^9^ observed a similar pattern in trout during sprinting, in which the trout activated their muscles earlier as they sprinted faster.

The earlier activation largely compensated for the increase in duty cycle, which means that the offset time for the muscle did not change significantly between steady swimming and acceleration (Table S4). This pattern is different than that for sprinting trout, in which the trout also shifted the offset time of their muscles earlier, keeping the duty cycle relatively constant^9^.

During accelerations, for midbody and posterior muscle, the change in phase corresponds to a large enough change in timing that the muscles likely produce substantial force during lengthening. During high accelerations, midbody and posterior red muscle are activated 49 ms and 43 ms before shortening begins. Red muscle in the pectoral fins of bluegill sunfish takes about 40-55 ms to reach peak force ^38^, based on twitch stimulation. Other fish species have similar durations for axial red muscle ^39^, although the properties may shift along the body ^14^. If axial red muscle in bluegill sunfish is similar, then it is likely that when the fish accelerates, this muscle generates substantial force during lengthening.

Moreover, during accelerations, muscles were not only active more often during lengthening (eccentric activation), but they also lengthened faster (Fig. 5C), particularly near the tail. Eccentric activity generally tends to lead to higher effective stiffness, and faster lengthening rates also increases stiffness ^34^.

**Figure 5.**
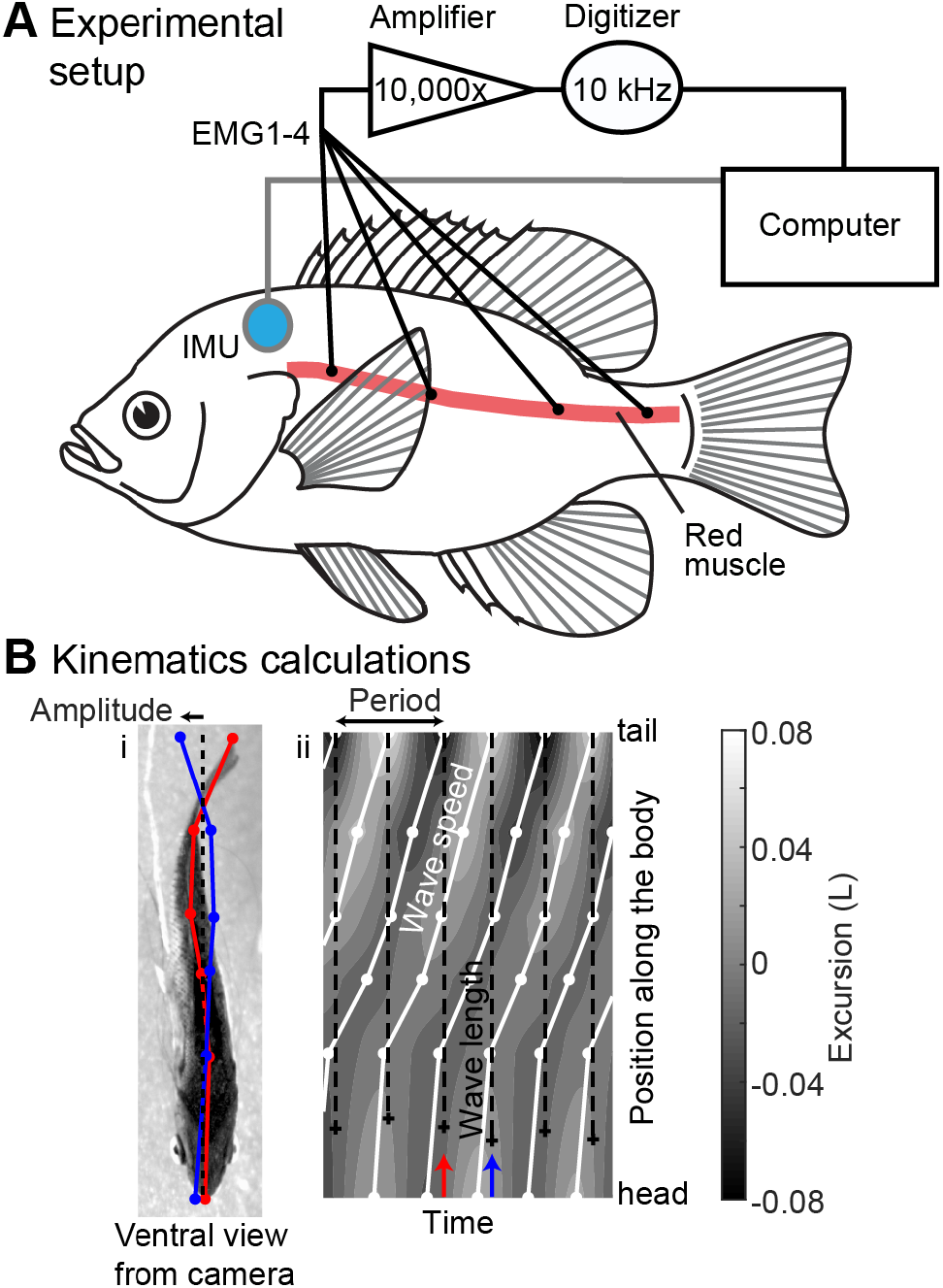
Experimental setup and swimming kinematics calculations. (A) Schematic representation of electromyographic apparatus used to record EMGs, showing longitudinal positions of the electrodes (four on each side), and the IMU apparatus used to record acceleration and its placement on fish. (B) Schematic of kinematics calculations. The left panel (i) shows one frame from ventral video, showing the digitized midline points (in red) and points from one half tail beat later (in blue). The right panel (ii) shows excursion of the body in gray, with peak excursion at each body position shown with connected white lines, and the body wavelength shown with a dashed line. Blue and red arrows indicate the time of the frames shown in panel B. See text for details of the calculations.

In escape responses, very rapid turning accelerations ^21^, duty cycle also increases, but primarily in the posterior region ^40^. Here, duty cycle and burst overlap also increased during accelerations, but more in the anterior region. Accelerations during escape responses are higher in magnitude (often > 3 *g*) ^21^ than even the high accelerations studied here (0.14-2.3 *g*, mean = 0.28 *g*). It may be that posterior duty cycle will increase more, similar to that in escape responses, during more rapid forward accelerations.

These measurements are consistent with the hypothesis that fish actively increase the stiffness of their bodies during accelerations. Long^7^ and Tytell^35^ stimulated both red and white muscle in eels and lampreys at different phases within a bending cycle and measured the resulting effective body stiffness. They both found that stiffness could more than double, and the maximum effective stiffness was achieved at an activation phase of -0.4 (using our phase convention; see Fig. 4A) and decreased as phase approached 0. If a similar phase relationship holds for the bluegill sunfish, then eccentric activity stiffens the tail most substantially, but also contributes to increasing effective stiffness in all body segments. Additionally, muscle stiffness in general increases during activity ^34^, and particularly during eccentric activity and as the strain rate increases ^41^. The mechanical effects of co-activation have not been studied in fishes, but work on humans showed that co-activation of trunk musculature increased trunk stiffness by 25 to 50% ^42,43^. Since the fish had the most bust overlap in anterior regions, co-activation likely contributes to stiffening the anterior body more than the tail, helping transfer the force from head to the tail to quickly propel the fish forward. Closer to the tail, eccentric activations at higher strain rates most likely increase stiffness to compensate for reactive forces from the environment, helping to stiffen the tail to transfer muscle force to the fluid ^8,20^ as the fish accelerates faster.

Based on our measurements, we propose that, in rapid movements like these accelerations, one of the important roles of red muscle is not only to produce power but to tune the effective body stiffness. Numerous studies, like ours, have found that red muscle is active during rapid movements when the contraction rate of the muscle is too low to provide useful power for the behaviour ^17, 44–46^. The ongoing red muscle activity may be a consequence of the motor neuron recruitment pattern for red and white muscle ^47^. In the accelerations here, cycle frequencies are almost always greater than 4 Hz. For pectoral fin muscles in bluegill sunfish, the power output begins to drop at frequencies above 3 Hz ^48^. Similarly, for largemouth bass (*Micropterus salmoides*), a closely related species, red muscle produces increasingly less power above frequencies of 3 Hz ^44^. Jayne and Lauder ^17^ proposed that force from red muscle helps to slow down the body’s movement and reverse the bending direction. This idea is part of our hypothesis. Because red muscle is more often active in lengthening during acceleration, it resists the body’s motion, which increases the effective stiffness. We suggest that one result of the ongoing red muscle activity during these rapid movements is to tune the effective body stiffness to increase overall performance.

This eccentric muscle activity may have an additional benefit: it does not require much metabolic energy. Higher forces are produced during eccentric activity ^49^ with very little energetic cost ^50,51^. Thus, by activating their muscle earlier, fish likely generate more force to stiffen the body against fluid forces, but without incurring large energetic costs. Similarly, in an *in vitro* test, a small amount of muscle co-activation increased total power output dramatically^52^.

However, eccentric muscle activity does not produce positive power for acceleration. The power for acceleration is likely coming from white muscle^17,40^. To accelerate in a fluid, bluegill sunfish require force both to accelerate their own mass, but also to overcome the acceleration reaction^53^, an additional force required to accelerate the fluid around them ^54^. Wise et al. showed that the acceleration reaction in bluegill sunfish is increased by the very same kinematics bluegill use during accelerations^53^. It may be that fishes can produce these additional forces economically by using eccentric activity.

Our results demonstrate one way that fish and other flexible organisms may approach the trade-off between steady swimming and rapid acceleration. To swim with low energy cost, computational modelling suggests that fish should minimize the energy required to bend their bodies ^3,5^; to accelerate rapidly, fish should have stiffer bodies ^5^. Measurements of passive body stiffness match this pattern. Fish that tend to swim slowly over long distances are much more flexible than those that are specialized for rapid acceleration^55^. Our results suggest that fish can tune their effective body stiffness in the same pattern, using red muscle activity to stiffen their bodies during acceleration.

## Materials and methods

### Animal care

Five adult bluegill sunfish (*Lepomis macrochirus* Rafinesque) were captured by beach seine in White Pond, Concord, MA, USA. All animals were housed individually in 38 l aquaria with 12 L:12 D cycle and were fed live worms or flake food daily. Water temperature (20±2°C) and pH (7.4) were kept constant and were equal to that used during experiments. Fish total length ranged from 14.5-16.5 cm (mean ± s.d. = 15.5±0.9 cm) and mass ranged from 42.0 to 66.0 g (57.0±9.7 g). Animal care and all experimental procedures followed protocols approved by Tufts University Institutional Animal Care and Use Committee (under protocols M2012-145 and M2015-149).

### Inertial measurement unit construction

An inertial measurement unit (IMU) was attached to each fish to collect rapid and accurate acceleration data (see additional sections below). Each IMU was constructed by soldering 1 m long coated, fine copper wire (80 μm diameter) to individual pads on a nine-axis (gyro, accelerometer, and compass) solid-state IMU chip (MPU-9250, InvenSense Inc.). Wires were soldered following the company’s instruction for the serial peripheral interface (SPI). The long copper wires were glued together with silicone to form a single cable and the IMU was waterproofed by applying several coats of epoxy (CircuitWorks Epoxy Overcoat, Chemtronics). Small wire loops were also epoxied to the IMU so it could be sutured onto a fish. Data was collected from the IMU by connecting it to a USB SPI interface (USB-8451, National Instruments) and data was recorded using a custom LabVIEW program (v. 2014, National Instruments).

### Electromyography and IMU attachment

Fine wire electromyography (EMG) electrodes were constructed by stripping <1 mm of insulation off the ends of a 1 m long double stranded, 55μm diameter steel wire and bending the bare ends into a hook. The wires were threaded through 27 gauge hypodermic needle for implantation.

Fish were anesthetized with a buffered 0.02% solution of tricaine methane sulfonate (MS222) for approximately 15 minutes and were weighed and measured. During surgery (< 2 h), anaesthesia was maintained by pumping buffered 0.01% MS222 over the fish’s gills. The electrodes were implanted into superficial, axial red muscle at four positions on both sides of the body in each fish (Fig. 5A). EMGs were positioned in the red axial muscle based on the following landmarks: EMG1 = fourth dorsal fin spine (40% of total body length), EMG2 = anterior edge of anal fin (53%), EMG3 = caudal edge of dorsal and anal fins (63%) and EMG4 = mid peduncle (74%) (Fig. 5A). Following electrode implantation, electrodes were secured by suturing the wires together to the fish’s skin. Each electrode had sufficient slack so that swimming movements did not remove the electrodes.

The IMU was sutured near the centre of mass on either the left or right side of each fish using three to four sutures (Fig. 5A). All EMG electrodes were glued together to form a cable that was sutured, along with the IMU cable, immediately anterior to the dorsal fin.

Each fish was allowed to recover for at least two hours after surgery. After completion of the experiment, fish were euthanized using an overdose of MS222 and were dissected to confirm that the electrodes were implanted in red muscle. EMGs implanted into white muscle were not included in the analysis since this muscle type is not active at slower speeds.

EMG signals were amplified 10,000 times using a differential AC amplifier (Model 1700, A-M Systems) and sampled at 10 kHz using a Powerlab digitizer (Model 16/35, A/D Instruments) and LabChart 7 software (v. 7.3.7, A/D Instruments). Signals were initially filtered with a 60 Hz notch filter, along with a band pass filter between 100 Hz and 1,000 or 5,000 Hz in the amplifier. Two 30 cm long copper rods and a 30×30 cm sheet of fine copper mesh were placed outside of the confined working section in the flow tank (described below) and were connected to a common ground to further reduce noise in EMG signals.

### Experimental procedure

Fish were placed individually in a 293 l recirculating flow tank (Loligo Systems) and were confined to a 90 cm long section of the 25×26×150 cm (height × width × length) working section. A high speed camera (Phantom Miro M120, Vision Research) filming at 200 frames per second was positioned underneath the flow tank to provide a ventral view of each fish within the working section. The camera, EMGs, and IMU were synchronized with a manual trigger.

For each of five fish, at least three steady swimming and three acceleration trials at several flow speeds (1.5, 1.75, 2.0, 2.25, 2.5 body lengths per second [L s^-1^]) were recorded. Each trial consisted of at least five tail beats. Fish swam in place during steady swimming trials by positioning rods in front of and behind the fish and ensuring that the fish did not drift in the video frame by more than 2 mm. In acceleration trials, fish were positioned in the rear portion of the confined section using rods and forward accelerations, in which the fish moved forward in the video frame, were elicited by moving the rods or by dropping an object behind the fish. Care was taken so that rods or falling objects did not come in contact with the fish. Fish accelerated forward by swimming faster than the flow tank speed and using relatively symmetrical, high-frequency tail beats. Extreme behaviours such as startle responses (indicated by a C-shape bend) and burst-and-glide swimming (a rapid tail beat or two followed by a powerless glide) were excluded from the analysis.

### Processing of video data

From each video, six locations on the body of the fish (snout, between the pectoral fins, anus, anterior base of anal fin, peduncle, and caudal fin tip) were manually identified using custom procedures in Matlab (R2014b, Mathworks) (Fig. 5Bi). Because the curvature in the anterior body is relatively low, these six points produce a good estimate of curvature in the locations of the EMG electrodes.

A custom Matlab program estimated kinematic parameters, identified EMG bursts and synchronized them with the tail movements, and used the IMU recordings to estimate body orientation and dynamic acceleration.

The true centre of mass position was estimated using the mass per length measured by Tytell ^56^ and the main body axis was estimated by fitting a straight line to the *x* and *y* distances to the centre of mass and then smoothing over 2 s. This duration was chosen because it was longer than the tail beat period, in order to smooth out fluctuations due to the tail oscillations.

The amplitude of movement at each body point was estimated by identifying positive and negative peaks in the excursion perpendicular to the main body axis (Fig. 1B, right panel) and tracking them as they moved along the body (white lines in Fig. 1B). All calculations were performed relative to arc length along the body. By tracking a peak along the body, we measured its position as a function of time; the average derivative of that value is the wave speed *V* (Fig. 5Bii). Tail beat period *T* was computed by taking the time difference between two peaks in lateral excursion on the same side of the body. Finally, we estimated the average body wavelength *λ* by multiplying the wave speed and the tail beat period:*λ*=*VT*.

The data were normalized to body lengths (L; head and tail amplitude, body wavelength) and body lengths per second (L s^-1^; body wave speed) to account for differences in the size of fish used in the study.

Body curvature *κ* was computed as

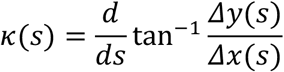

where *∆x* and *∆y* are the differences in *x* and *y* position of successive points along the midline, and *s* is the arc length along the midline.

### Processing of EMG data

Spikes were identified in EMG recordings based on a manual threshold, and bursts of spikes were identified by finding groups of spikes separated by less than a manually chosen interburst interval (0.05 s). Bursts with fewer than three spikes were excluded. Burst duration was the time between the first and last spike in a group and was normalized relative to the current cycle period to give the duty cycle (= proportion of strain cycle period). Burst overlap was the time between the beginning of a burst on one side and the end of the previous burst on the other side. A burst overlap greater than zero indicates co-activation, while a negative value indicates a gap between bursts.

The timing of maximal body curvature was interpolated to the specific position of each EMG electrode and a continuous curvature phase was estimated based on the timing of positive and negative peaks and positive- and negative-going zero crossings. This phase variable then gave us the onset and offset phase for each burst, relative to the local curvature.

Red muscle strain and strain rate were approximated using the local curvature, following Coughlin et al. ^57^. Strain *∊* and strain rate were *∊ ̇*computed as

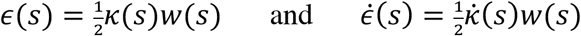

where *w* is the total width of the body at a position *s*. We then averaged the strain and strain rate during the EMG burst at that location, and we also separately averaged the lengthening and shortening rates during EMG bursts.

### Processing of IMU data

The IMU has a three-axis gyroscope, which measures angular velocity ***ω***, and a three-axis accelerometer, which measures total acceleration **a**=**a**_*dyn*_+**g**, where **a**_*dyn*_ is dynamic acceleration of the center of mass and **g** is gravitational acceleration. The sampling rate of the IMU was 200 Hz. To separate the dynamic acceleration, we needed to estimate the orientation **q** of the body, from which we can compute **g** and subtract it from the total acceleration.

First, the constant bias of the gyro was estimated by recording for 60 s when the sensor was motionless in the water. The mean bias was then subtracted from further recordings. The gyro signal was then filtered with a bandpass filter between 0.5 and 10 Hz.

We used the algorithm from Madgwick and colleagues (2011) to estimate orientation. Briefly, orientation can be computed by integrating the angular velocities or, when **a**_*dyn*_ is low, **a** can be used as a good estimate of **g**. We first integrated angular velocities using quaternions to avoid singularities with Euler angles. The quaternion angular velocity 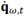 based on the gyroscope at the current time *t* was defined based on the previous estimate of the orientation 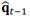,

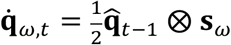

where *s_ω_*=[0 *ω_x_ ω_y_ ω_z_*]. Then the angular velocity was updated based on a gradient descent toward the orientation defined by the accelerometer reading

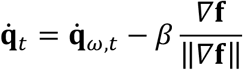

where

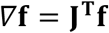

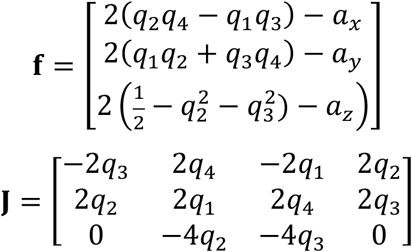

and 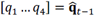 are the four elements of the previous quaternion estimate of the orientation and [*a_x_ a_y_ a_z_*]=**a** are the components of the current accelerometer reading. We used *β*= 2.86 deg s^-1^, the optimal value based on the maximum angular velocities we measured ^58^.

Then we updated the previous orientation estimate

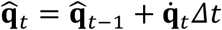

which gave an estimate of the gravity vector in the sensor’s coordinate system

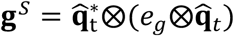

where *e_g_*=[0 0 0 –1] is the quaternion gravitational vector in the world coordinate system and **q**^*^ denotes the conjugate of **q** The dynamic acceleration is thus **a**_*dyn*_=**a**_*tot*_–**g**^s^. The initial quaternion orientation was estimated based on the mean accelerometer signal in the first half second, **ā** _*init*_=[*a_x_*_,*init*_ *a_y_*_,*init*_ *a_z_*_,*init*_],

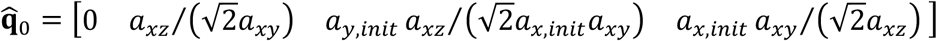

where 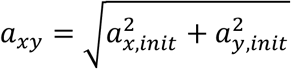 and 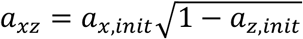

We calibrated the orientation of the sensor on the fish by measuring the gravitational vector while holding the fish on its side, with its snout downwards, and in a normal dorsal-side up position, as the fish was reviving from the anaesthesia. We then constructed a rotation matrix from the sensor’s coordinate system to the fish’s, using the Gram-Schmidt method ^59^ to ensure that the fish’s coordinate system had orthogonal basis vectors.

Once the dynamic acceleration was estimated, we took the peak value in the fish’s forward direction during each half tail beat (when the tail swept from left to right or the other way). We selected peak acceleration since this value most clearly synchronized with the tail beats.

### Data analysis

All statistical analyses were performed in R version 3.4.4^60^. Regressions were estimated using the nlme package, version 3.1-131.1 ^61^ and marginal means were estimated using the emmeans package, version 1.2.1 ^62^. Figures were created using ggplot2, version 2.2.1 ^63^.

For the kinematic data, we first grouped the acceleration into four trial categories based on the fish’s forward acceleration measured by the IMU (steady; low: *a_dyn_* < 0.07 *g*; medium: 0.07 *g* ≤ *a*_dyn_ < 0.14 *g*; high:*a*_dyn_ ≥ 0.14 *g*). Acceleration was binned because it had a strongly non-normal distribution, which would have biased standard regression results. Instead, we grouped the measurements into bins so that we had approximately the same number of unsteady acceleration tailbeats in each bin. This procedure is similar to the ranking procedures that are the basis of most nonparametric statistics^64^. We tested the robustness of the ANOVA models below by varying the number of bins and the methods for choosing them. None of the overall patterns or statistical results changed.

A two-way mixed ANOVA model with autoregressive covariance structure (AR1) was performed for each kinematic variable (tail beat frequency, body wave speed, head and tail amplitudes, body wavelength) to determine any significant relationships between acceleration categories, swimming speed, and the interaction between acceleration and swimming speed (=fixed effects), while accounting for individual variations among the different trials (=random effect) and sequential tail beats (=repeated effect). The autoregressive structure was used because sequential tail beats are not statistically independent of one another. We also performed a principal components analysis followed by a multivariate ANOVA^65^ to determine whether the swimming kinematics as a whole changed due to acceleration, swimming speed, and their interaction.

For the EMG data, a similar three-way mixed ANOVA model with autoregressive covariance structure (AR1) was performed for each variable (burst duration, duty cycle, burst overlap, strain rate during active muscle shortening and lengthening) to determine any significant relationships between acceleration, EMG position, swimming speed, and all of their interactions (=fixed effects) while accounting for individual variations among the different trials (=random effect) and sequential tail beats (=repeated effects). Two-way circular ANOVA tests (Harrison-Kanji tests) were performed using the CircStat Toolbox ^27^ in Matlab (R2017b, Mathworks, Inc., Natick, MA) to assess correlations between the onset of EMG bursts and peak acceleration by EMG location.

All mean values are reported as mean±s.d. Because of the large number of statistical tests, we use a significance cutoff of p < 0.01 rather than 0.05.

## Data and code accessibility

The data and code that support the findings of this study are available at doi: 10.25833/ywtf-9c79.

## Acknowledgements

We thank Vishesh Vikas for assistance with the IMUs, Nicole Danos for discussions about this manuscript, and Tytell lab members for assistance with animal husbandry. This work was supported by the US Army Research Laboratory and the US Army Research Office under Contract/Grant Nos. W911NF-14-1-0494 and W911NF-14-1-0268 (to E.D.T.), the National Science Foundation under grant RCN-PLS 1062052 (to Lisa J. Fauci and Avis H. Cohen) and the National Institute of Health Training in Education and Critical Research Skills under grant number K12GM074869 (to M.A.B.S).

## Author Contributions (names as initials)

MABS and EDT designed the experiments. MABS, TNW, and ALB performed the experiments and digitized the videos. MABS and EDT analysed the data and wrote the manuscript. All authors reviewed the manuscript.

